# Snow is an important control of plant community functional composition

**DOI:** 10.1101/564583

**Authors:** Konsta Happonen, Juha Aalto, Julia Kemppinen, Pekka Niittynen, Anna-Maria Virkkala, Miska Luoto

## Abstract

The functional composition of plant communities is a critical modulator of climate change impacts on ecosystems, but it is not a simple function of regional climate. In the Arctic tundra, where climate change is proceeding the most rapidly, communities have not shifted their trait composition as predicted by spatial temperature-trait relationships. Important causal pathways are thus missing from models of trait composition change. Here, we study causes of plant community functional variation in a tundra landscape. We consider the community weighted means of plant vegetative height, as well as two traits related to the leaf economic spectrum. Specifically, we model their responses to locally measured summer air temperature, snow conditions, and soil resource levels. For each of the traits we also quantify the importance of intraspecific trait variation (ITV) for between-community functional differences and trait-environment matching. Our study shows that in a tundra landscape 1) snow is the most influential abiotic variable affecting functional composition, 2) vegetation height is under weak local environmental control, whereas leaf economics is under strong local environmental control, 3) the relative magnitude of ITV differs between traits, and 4) ITV is not very consequential for community-level trait-environment relationships.

## Introduction

Contemporary climate change in the Arctic is so rapid that the ecosystem changes it causes can be observed over ecological timescales (Post et al. 2009). While warming itself will alter the flows of matter and energy through ecosystems, these changes can be heavily modulated by changes in the functional composition of plant communities (Díaz et al. 2004). Recent research has shown that above-ground trait variation between plant species and communities can be compressed into two independent trait axes: plant size and the leaf economics spectrum (Díaz et al. 2015; Bruelheide et al. 2018). Empirical evidence has shown how the functional composition of Arctic tundra communities along these axes has recently changed (Bjorkman et al. 2018a), however we still do not understand the environmental drivers behind this development. Such inability to explain and model trait composition and its spatio-temporal dynamics reflects the fundamental gaps in our understanding of the drivers of plant community functional composition (Bruelheide et al. 2018).

In any given landscape, the trait variation among species pools and communities can cover most of the global trait spectrum, especially for traits related to leaf economics (Wright et al. 2004; Bruelheide et al. 2018). This suggests that environmental factors with large variation at local to landscape scales, such as soil resources and disturbance, may be more important determinants of plant community trait composition compared to macroclimatic gradients (e.g. mean air temperature and/or precipitation) (Bruelheide et al. 2018). One such factor, which has received limited attention in trait-environment studies, is snow. The importance of snow for vegetation in cold climates has long been recognized in vegetation research, yet often ignored (Niittynen et al. 2018). Snow can influence optimal trait values by various mechanisms, such as protecting vegetation from extreme cold temperatures, frost, and abrasion by wind-blown ice particles, as well as providing water sometimes long into the summer, and limiting the length of the growing season (Braun-Blanquet et al. 1932; Niittynen et al. 2018). Its effects on functional composition in the modern sense have been less commonly studied, particularly in combination with other key environmental variables.

It is increasingly recognized that intraspecific trait variation (ITV) is both responsible for a large fraction of trait variation between communities, and important for the outcomes of ecological processes (Siefert et al. 2015). However, further evidence is needed on whether this variation has an effect on community trait-environment relationships. This is because even if ITV is large and adaptive at the species level, species-specific responses do not necessarily scale up to the community level (Lajoie and Vellend 2015), in which case between-community differences in functional composition caused by ITV would be random.

Here we study the causes of variation in functional composition of plant communities in a tundra landscape. Our study focuses on the community weighted means (the functional markers in Garnier et al. 2004) of three traits that represent the two above-ground trait axes: vegetative height, specific leaf area (SLA), and leaf dry matter content (LDMC). We partition between-community trait variation to contributions from species turnover and intraspecific effects. To examine the environmental controls of community functional composition, we model the responses of community weighted mean traits over three environmental gradients: temperature, soil resources (moisture and pH), and maximum snow depth. To test if intraspecific variation in these traits affects trait-environment relationships, we examine how aggregating the species trait data at global, landscape, and local resolutions affects the strength of the trait-environment relationship.

## Methods

### Study site and abiotic variables

The study area is located in Kilpisjärvi, northwestern Finnish Lapland (N69.06°, E20.81°) with a mean annual air temperature of −1.9°C and annual precipitation sum of 487 mm (1981-2010; Pirinen et al. 2012). The area is part of the oroarctic mountain tundra, characterized by a high abundance of seasonal snowbeds and steep environmental gradients related to elevation and mesotopography (Virtanen et al. 2016). Vegetation at the study site is a mosaic of dwarf-shrub heaths and meadows (see list of observed species and their abundances in Appendix S1: Table S1). Plants are grazed by reindeer during the growing season, and by voles and lemmings throughout the year.

In summer 2016, a 1.5 km by 2 km environmental monitoring network was established in the tundra between the mountains Saana and Jehkas, at 580–920 meters above sea level. A schematic map of the study area and the different environmental measurement schemes is given in Appendix S1: Figure S2.

Soil moisture was measured during the growing seasons of 2016–2018 at 220 locations, 3–6 times per growing season. In most cases, moisture measurements in each location were done at least 24 hours after any rainfall event. Three measurements were taken and averaged in five 1 m^2^ plots: one central plot and one plot 5 meters away in each cardinal direction. Measurements were taken using a hand-held time-domain reflectometry sensor (FieldScout TDR 300; Spectrum Technologies Inc., Plainfield, IL, USA). Snow depth was measured with a metal rod in early April of 2017 and 2018 at the time of maximum snow depth, from the same five plots of the moisture measurement scheme. Temperature loggers (Thermochron iButton DS1921G and DS1922L; temperature range between −40°C and 85°C, resolution of 0.5°C, and accuracy of 0.5°C) were installed in 112 locations to monitor temperatures 10 cm above and below ground, at 2–4 hour intervals. These single time-point measurements were consequently aggregated to monthly averages.

Samples of the organic soil layer were collected from 200 locations in August of 2016 and 2017. Soil samples were freeze-dried following SFS 3008, and pH was analyzed from the samples in the laboratory of the University of Helsinki, following ISO standard 10390. The soil samples were taken ca. two meters away from the central plot of the moisture measurement scheme to avoid perturbation. The distance between two adjacent locations was a minimum of 23 meters (average 101 meters). Individual study locations were thus in different vegetation patches.

We calculated average July air temperatures, growing season soil moisture levels, and maximum snow depths for each location with random effects models using the R package lme4 (version 1.1-18-1, Bates et al. 2015). Random effect models allow the use of hierarchical structure in the data to pool information over those hierarchies. Thus calculated averages are more robust to outlier observations (Bates et al. 2015). Temperatures were modelled using location and year as random effects (*y* ∼ (1|*location* + *year*)). Snow depth and soil moisture were modelled using plot nested in location, and year as random effects (*y* ∼ (1|*location*/*plot* + *year*)). Soil moisture was log_e_-transformed before modelling. Finally, the environmental variables were predicted for the location (temperature), or for the central plot of the moisture measurement scheme (snow depth and soil moisture). Soil moisture was back-transformed before further analyses.

The average soil moisture and pH of the organic layer were highly correlated (*ρ* > 0.8). Hence, we reduced them to their first principal component, which we hereafter refer to as the soil resource axis. This axis accounts for over 83% of the overall variability.

### Species composition and functional traits

We quantified vascular plant community composition in 143 locations with the point intercept method, using a 20 cm diameter circular frame with 20 evenly spaced pinholes. An image of the frame can be found in Appendix S1: Figure S3. The frame was placed as close to the environmental measurement plot as possible without disturbing the measurements (on average, 2.9 meters away). We measured the abundance of each species as the total number of times that species touched pins (3 mm in thickness) lowered into the vegetation through the frame. The three traits in our landscape were also measured within the point intercept frame. For each species that touched a pin in a plot, the height of the highest leaf was measured on two randomly selected individuals. Two leaf samples per species, from different individuals where possible, were also taken and stored in a resealable plastic bag with moist paper towels. After each field day, the leaf samples were stored at 4°C for up to three days until further processing.

We weighed leaf samples for fresh mass using a precision scale, and scanned them at a resolution of 600 dpi using a document scanner. The samples were then dried at 70°C for 48 hours and weighed for dry mass. We measured leaf area from the scanned leaf images using the Fiji-distribution of the ImageJ software (version 1.52h, Schneider et al. 2012; Schindelin et al. 2012). Using these measurements, we calculated SLA (area/dry weight; mm^2^/mg) and LDMC (dry weight/fresh weight; unitless). We downloaded global observations on vegetative height, SLA, and LDMC for our study species from the Tundra Trait Team (TTT) database (Bjorkman et al. 2018b).

We calculated community weighted mean (CWM) values for the three traits at three resolutions. CWM_local_ were calculated with traits measured in one study plot only. When calculating CWM_landscape_, we used one trait value – the average of all observations in the landscape – for each species. CWM_global_ was calculated using trait values from the TTT database. CWMs were calculated for a community only if trait data was available for the species that formed over 90% of the total abundance of the community. All CWMs were log_e_-transformed before analyses.

### Statistical analyses

All analyses were performed in R version 3.4.4 (R Core Team 2018).

We quantified the contributions of species turnover and ITV to CWM_local_ in all 143 surveyed plots using the method reported in Lepš et al. (2011). Briefly, the method is based on partitioning the variance of CWM_local_ to effects from CWM_landscape_ and ITV (CWM_landscape_ - CWM_local_). Calculations were done with the function varpart in the R package vegan (version 2.5-2, Oksanen et al. 2018).

We modelled the responses of CWM_global_, CWM_landscape_, and CWM_local_ to July mean temperature, soil resources, and snow depth using version 1.8-27 of the mgcv package for generalized additive models (GAMs) (Wood 2011). For each model, we used the maximum amount of observations possible. Due to different coverage of species in the local, landscape, and global resolution trait data sets, the final number of observations varied between 87 and 94. We set the basis dimension of our smooth terms to three in order to restrict model overfitting. In our experience, this allows enough freedom for the model to represent realistic ecological responses. We evaluated the predictive power of our models with leave-one-out cross-validation. That is, we fitted each models N additional times, each time leaving out one observation, and used the model to predict that left-out observation. We summarized the predictive power of each model as the squared correlation of observed and predicted CWMs.

The fitted CWM in a location is the sum of the modelled intercept and smooth terms in its environmental conditions. To investigate which variables made the greates contribution to explaining the CWMs, we calculated the unique contribution of each environmental variable to multiple R^2^ by subtracting its smooth term from the fitted values while keeping the other smooth terms constant. The resulting increase in error variance is the unique contribution of that smooth term to the explained variation. To allow direct comparison of the predictors’ relative contributions, we standardized the values to total 1.

## Results

The relative magnitude of ITV differed between traits (Figure 1 a). Most of the variation (62%) in community mean height was due to intraspecific variation, while variation in leaf traits was more due to species turnover (83% and 47% of total variation for SLA and LDMC, respectively). Turnover and intraspecific effects were correlated for LDMC. Thus, a large fraction of its variation could not be uniquely partitioned (the joint partition in Figure 1 a).

**Figure 1:**
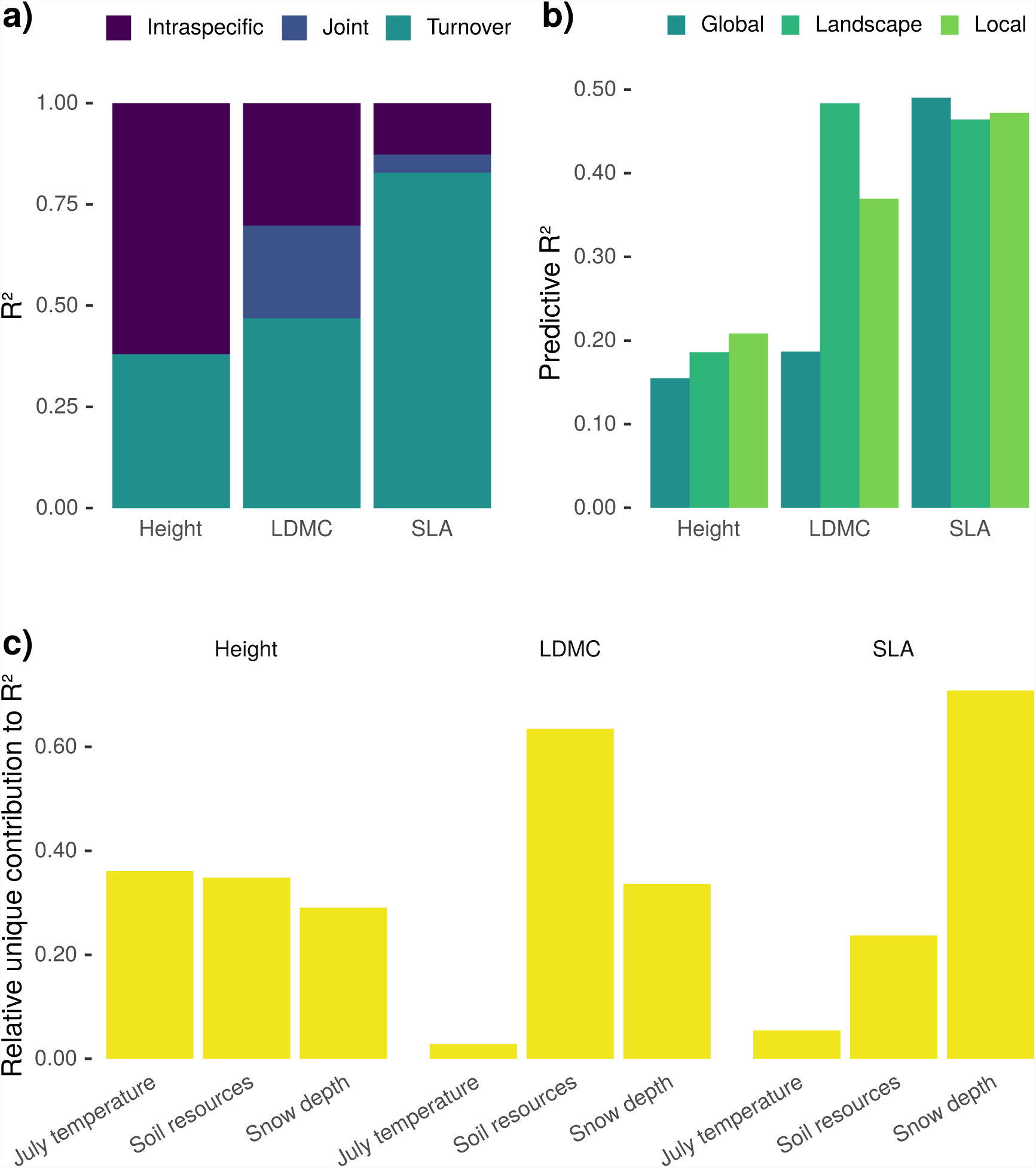
a) Between-community functional differences decomposed to contributions from ITV and species turnover. Joint contributions reflect correlations in the direction of intraspecific and turnover effects on community functional composition. b) Squared correlation of leave-one-out predicted and observed community weighted mean trait values. Results are from GAMs fitted with three different resolutions of species trait data: CWM_global_, CWM_landscape_, and CWM_local_. c) Relative unique contribution of each environmental variable to multiple R^2^ of models explaining CWM_landscape_.

The cross-validation of GAMs in explaining trait CWMs showed that traits differed significantly in how well they could be predicted from environmental conditions. Height models attained the lowest (∼ 0.20) and SLA models the highest (∼ 0.45) predictive performances (Figure 1 b). Predictive performance was dependent on trait measurement resolution. Different traits had different optimal resolutions, but, on average, models calibrated using landscape resolution data attained the highest predictive performance. This was due to the LDMC models, whose predictive performance was significantly worse for both local and global resolution data.

For brevity, the following results consider CWM_landscape_ only. The results for other trait measurement resolutions, which are qualitatively similar, are presented in supplemental figures.

Each trait had a different environmental variable as the most influential predictor (Figure 1 c). July air temperature had the highest contribution to the explained variance in community mean height, soil resources in LDMC, and snow depth in SLA. Variable importance for models using CWM_local_ and CWM_global_ are presented in Appendix S1: Figure S4.

The smooth terms and their confidence intervals from GAMs explaining CWM_landscape_ are presented in Figure 2. Height had a positive linear response to July temperature, and a negative saturating response to soil resources. The response of height to snow was negligible at maximum snow depths of up to 100 cm, and negative at deeper snow depths. SLA had a positive linear relationship with temperature (although the confidence interval overlapped zero), a positive linear response to soil resources, and a unimodal response to snow depth. The responses of LDMC to the environmental variables were opposite to those of SLA. The modelled responses for CWM_local_ and CWM_global_ are presented in Appendix S1: Figure S5.

**Figure 2:**
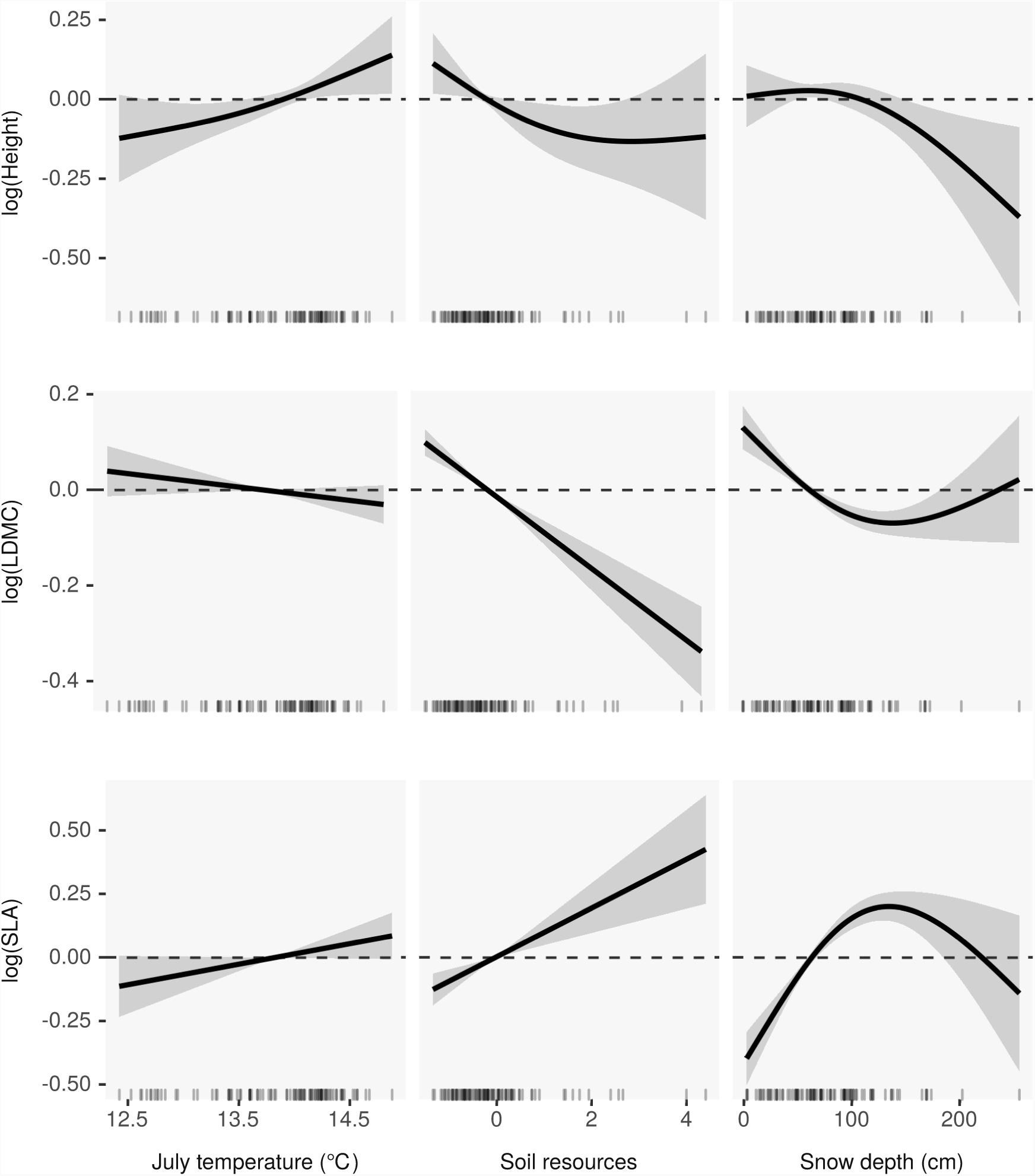
Thin plate splines and their confidence intervals (2×SE) describing the effects of three environmental variables on community weighted mean trait values in a treeless tundra landscape. At this scale the effects of soil resources and snow on functional composition in these communities are stronger than the effect of summer temperature. The rugplots at the bottom edge of each subplot show the distributions of each explanatory variable.

## Discussion

Intraspecific trait variation (ITV) was responsible for a higher proportion of between-community functional differences in height compared to leaf economic traits. In our study area, reindeer grazing reduces the height and cover of tall, deciduous shrubs, such as willows (*Salix* spp.) and dwarf birch (*Betula nana*) (Pajunen et al. 2008). Temperatures are also quite harsh, especially for plants protruding above the snow during the winter. These characteristics may severely constrict the height variation between species. Moreover, factors with strong effects on leaf economics are not similarly limited in the area. As a result, the communities in our study are likely to represent a sizable portion of global variation in community averaged leaf economic trait values. Many studies reporting high ITV contributions to community functional turnover have focused on clearly defined vegetation types with, perhaps, low variation in environmental factors that have strong effects on traits (table S3 in Siefert et al. 2015). This would cause contraction of viable species trait values and their CWMs, hence inflating the importance of ITV (Auger and Shipley 2013). Our interpretation is that the high proportion of intraspecific variation in height is caused by the contraction of interspecific height differences owing to limitation by temperature and grazing.

Around 23% of the between-community variation in LDMC could not be attributed to intraspecific and species turnover effects. This indicates that the effects are correlated: locations that harbor species with low LDMC also have individuals with lower LDMC than expected for their species. This may be due to water availability, which both promotes species with lower LDMC, and is the mechanistically linked to the leaf water content of plant individuals (Gardner 1965).

Accounting for ITV, especially within the studied landscape, was not essential for predicting community functional composition from environmental conditions. Height variation between communities was relatively low (CWMs between 1–20 cm, Appendix S1: Figure S6). In addition, age differences between individuals of up to 150 years (Büntgen et al. 2015) cause non-adaptive ITV in height. Thus, explaining the variation in community mean height is challenging and results in low predictive power. Yet, landscape and local resolution trait measurements seemed to provide more information on height-environment linkages. Our results are in line with the global analysis by Bjorkman et al. (2018a), which demonstrated that species across the tundra have significant positive intraspecific height responses to air temperature.

SLA was relatively well predicted by environmental conditions, and finer resolution trait measurements did not produce stronger trait-environment relationships. This may be explained by the minor difference between CWMs calculated at different trait resolutions (Appendix S1: Figure S6), which suggests that ITV in SLA has little impact on community-level trait-environment matching across the tundra.

Interestingly, LDMC-environment correlations were maximized when the model was fitted to the intermediate resolution trait data. Changing the resolution of trait measurements from landscape to local or global decreased the predictive power significantly. LDMC is generally highly correlated with SLA, and in our data landscape resolution LDMC was more correlated with SLA than local LDMC (Appendix S1: Figure S6). ITV in LDMC apparently consists of plastic responses to local water availability, while environmental filtering seems to operate on something which is more accurately captured by SLA and the landscape-level average LDMC.

Vegetation height was weakly explained by temperature, soil resources, and snow. In addition to a positive temperature effect, our models identified a negative effect of deep snow on plant height. This is in line with previous research showing negative effects of snow-cover duration on vegetation height (Choler 2005; Venn et al. 2011). Curiously, higher soil moisture and soil pH levels appeared to cause decreases in vegetation height. This is in contrast to the predictions of resource competition theory (Tilman 1988). Since vegetative height in this landscape is strongly limited by herbivory and air temperature, increasing resource levels do not necessarily lead to increased plant height (Kaarlejärvi et al. 2013). Decreasing height along the soil moisture and pH gradient is probably due to high resource levels favoring short-statured, grazing resilient herbaceous species (Eskelinen 2008).

Leaf economic traits were strongly controlled by soil resources and snow. Resource availability is a major control of leaf economics, which is well established in the literature (eg. Garnier et al. 2004; Pérez-Ramos et al. 2012; Spasojevic and Suding 2012). Our results support the positive linear response of community leaf economics to resource availability. However, the effects of snow on community position along the leaf economics spectrum is not as well studied. We found a unimodal effect of snow depth on leaf economic traits, such that the highest SLA and lowest LDMC were observed at snow depths of roughly 120 cm. Incidentally, this is also the depth where protection against low temperatures by the snowpack seems to saturate (Appendix S1: Figure S7). Choler (2005) observed a positive correlation between late snow melting day and higher SLA. There is a high correlation between landscape scale snow depth, growing season length, and plant perceived temperatures during the coldest time of the year (Appendix S1: Figure S7). Therefore, we cannot partition the snow depth-leaf economics relationship into effects from protection and shortening of the growing season. However, the unimodal response observed in our data suggests that there might be a switch in the driving process when snowpacks reach their maximal protective capacity, such that plants perceive any additional snow as stress rather than protective.

The links between the changes in snow cover, snow depth, and temperatures in Arctic systems are not simple (Niittynen et al. 2018). While the duration of snow cover has declined by an average of two to four days per decade, the rate of change varies across the Arctic (AMAP 2017). Reduction in sea-ice cover causes increased precipitation in the Northern hemisphere, which may even increase snowfall and maximum snow depth in many areas (Liu et al. 2012). This indicates that changes in snow depth are not tightly coupled to changes in snow cover duration and temperature across the Arctic. Experimental work and pan-Arctic comparative studies are needed to robustly study what influences the snow-trait relationships. As a first step, existing data from snow manipulation experiments could be re-analyzed in a functional trait framework.

Community leaf economics are strongly controlled by local scale factors that have complex relationships with macroclimate. It is thus not surprising that the observed changes in the position of tundra plant communities on the fast-slow continuum do not follow predictions made by regressing leaf traits against temperature alone, but instead seem to remain stable (Bjorkman et al. 2018a). The observed stasis could perhaps be due to warmer temperatures and diminishing snow causing opposing selection pressures on the leaf traits of plant communities.

## Acknowledgements

JA and ML were funded by the Academy of Finland (project numbers 307761 and 286950, respectively). AMV was funded by the Academy of Finland (project number 286950), Otto A. Malm foundation, Societas pro Fauna et Flora Fennica, Nordenskiöld-samfundet, The Finnish Cultural Foundation, Alfred Kordelin Foundation, and Väisälä fund. JK was funded by the Doctoral Programme in Geosciences of the University of Helsinki, Tiina and Antti Herlin Foundation, and Maa-ja vesitekniikan tuki ry. We thank Elina Kaarlejärvi for thoughtful comments on this manuscript, and Jacquelin DeFaveri for language revision.

