## Appendix S1 for "Snow is an important control of plant community functional composition"

### Supporting information for the article "Snow is an important control of plant community functional composition"

2019-03-01

**Appendix S1**

**Table S1.** List of species and their total point-intercept frequencies in the 143 surveyed plots. Nomenclature follows Tropicos (http://www.tropicos.org).

| Species | n |
|---|---|
| *Empetrum nigrum* | 1089 |
| *Vaccinium myrtillus* | 625 |
| *Betula nana* | 439 |
| *Deschampsia flexuosa* | 358 |
| *Phyllodoce caerulea* | 324 |
| *Vaccinium vitis-idaea* | 306 |
| *Carex bigelowii* | 132 |
| *Salix herbacea* | 75 |
| *Juniperus communis* | 70 |
| *Juncus trifidus* | 67 |
| *Nardus stricta* | 53 |
| *Festuca ovina* | 47 |
| *Vaccinium uliginosum* | 45 |
| *Cassiope tetragona* | 43 |
| *Cassiope hypnoides* | 36 |
| *Linnaea borealis* | 33 |
| *Loiseleuria procumbens* | 29 |
| *Lycopodium annotinum* | 26 |
| *Bistorta vivipara* | 24 |

| | |
|---|---|
| *Carex vaginata* | 18 |
| *Solidago virgaurea* | 18 |
| *Viola biflora* | 18 |
| *Antennaria alpina* | 14 |
| *Calamagrostis stricta* | 14 |
| *Carex dioica* | 14 |
| *Antennaria dioica* | 13 |
| *Anthoxanthum odoratum* | 12 |
| *Dryas octopetala* | 12 |
| *Sibbaldia procumbens* | 12 |
| *Carex lachenalii* | 9 |
| *Pedicularis lapponica* | 9 |
| *Carex brunnescens* | 8 |
| *Taraxacum* sp. | 8 |
| *Trientalis europaea* | 8 |
| *Trollius europaeus* | 8 |
| *Alchemilla* sp. | 7 |
| *Cornus suecica* | 6 |
| *Andromeda polifolia* | 5 |
| *Carex magellanica* | 5 |
| *Diapensia lapponica* | 5 |
| *Thalictrum alpinum* | 5 |
| *Diphasiastrum alpinum* | 4 |

| | |
|---|---|
| *Selaginella selaginoides* | 4 |
| *Agrostis mertensii* | 3 |
| *Bartsia alpina* | 3 |
| *Campanula rotundifolia* | 3 |
| *Eriophorum angustifolium* | 3 |
| *Gnaphalium supinum* | 3 |
| *Leontodon autumnalis* | 3 |
| *Trisetum spicatum* | 3 |
| *Carex bigelowii x aquatilis* | 2 |
| *Hieracium alpinum* | 2 |
| *Minuartia biflora* | 2 |
| *Poa alpina* | 2 |
| *Potentilla palustris* | 2 |
| *Salix glauca* | 2 |
| *Salix lapponum* | 2 |
| *Salix polaris* | 2 |
| *Arctostaphylos alpina* | 1 |
| *Astragalus alpinus* | 1 |
| *Calamagrostis lapponica* | 1 |
| *Carex rotundata* | 1 |
| *Equisetum sylvaticum* | 1 |
| *Juncus biglumis* | 1 |
| *Petasites frigidus* | 1 |

| | |
|---|---|
| *Potentilla crantzii* | 1 |
| *Ranunculus nivalis* | 1 |
| *Rubus chamaemorus* | 1 |
| *Saussurea alpina* | 1 |
| *Veronica alpina* | 1 |

**Figure S2.** A schematic map of the study area and the different environmental measurement schemes. The inset describes how the different measurement plots are positioned within one location. Aerial photographs from the National Land Survey of Finland (CC-BY).

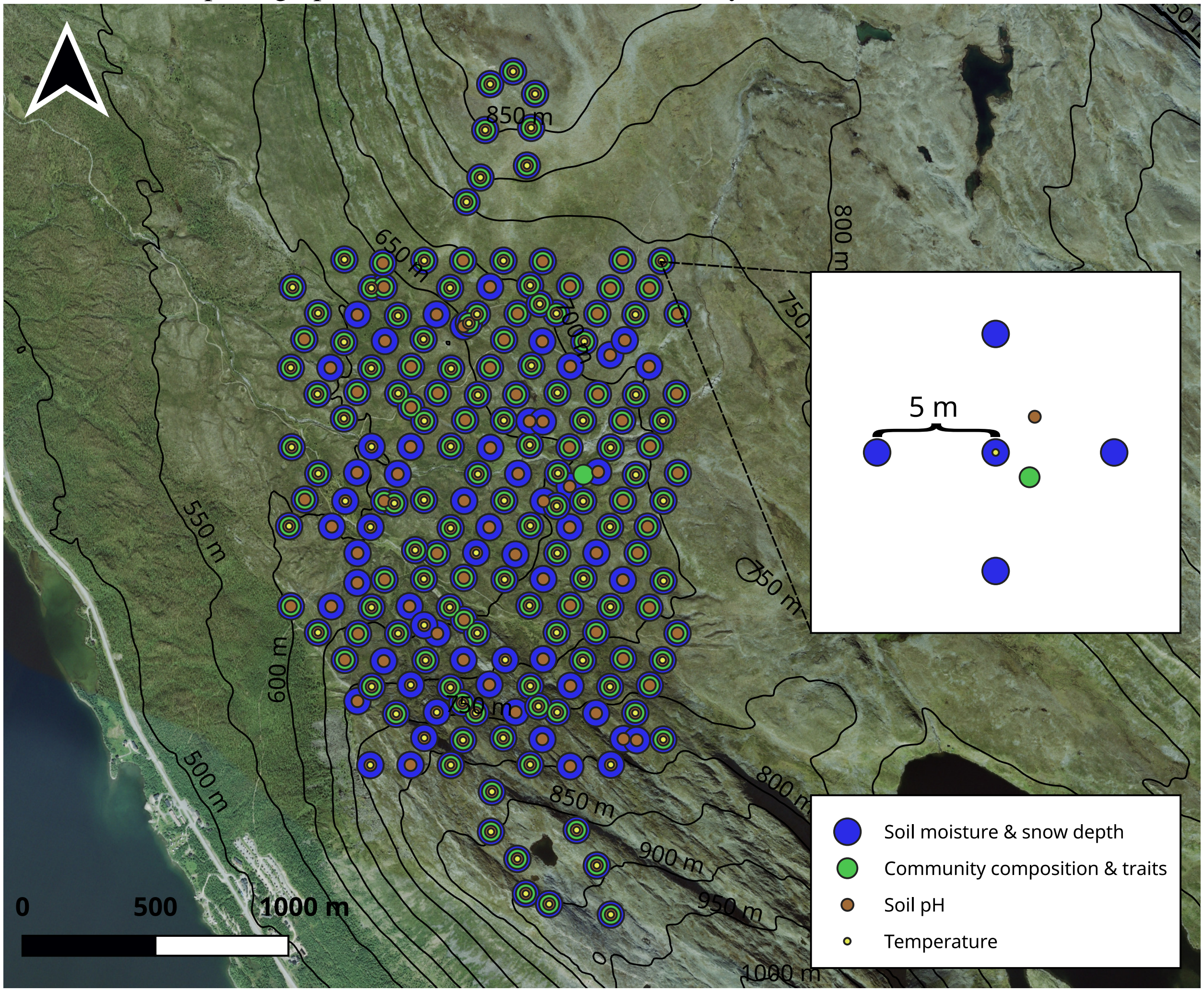

**Figure S3.** The point-intercept frame used in this study.

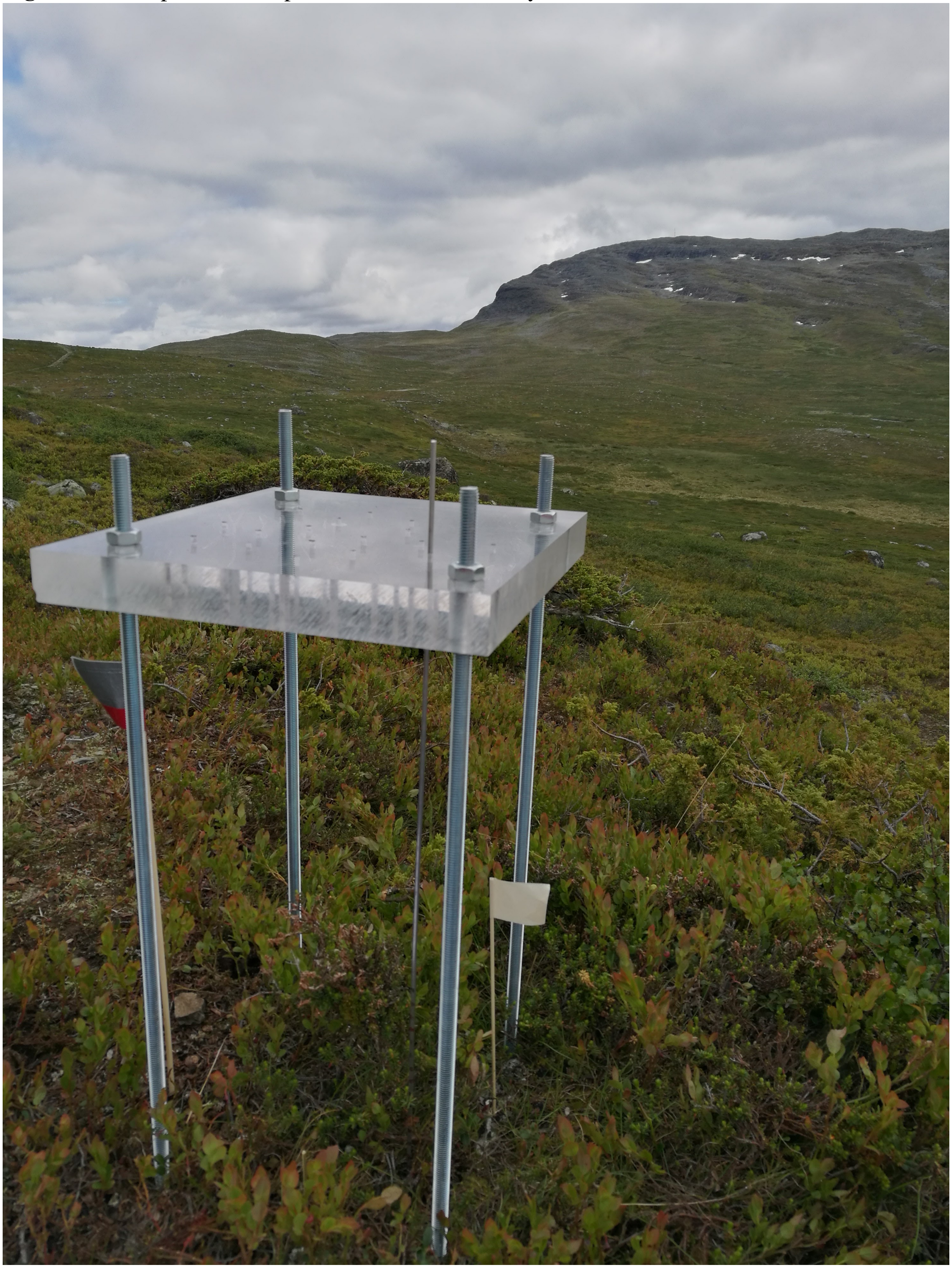

**Figure S4.** The relative unique contribution of July mean temperature, soil resources and maximum snow depth for explaining variance in $CWM_{local}$, $CWM_{landscape}$, and $CWM_{global}$ for height, LDMC and SLA.

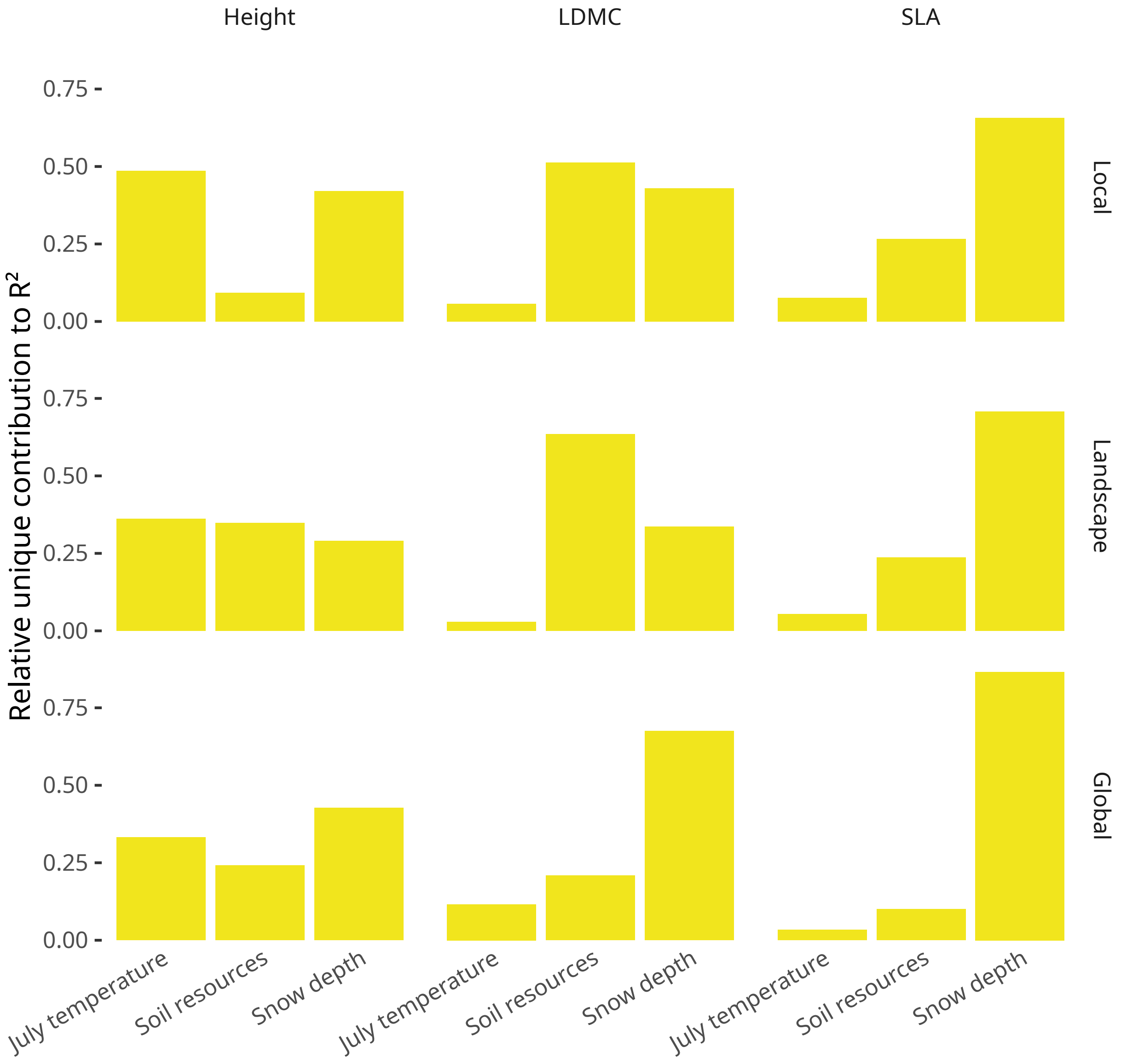

**Figure S5.** Smooth terms and their confidence intervals (SE*2) from models of $\log_e$-transformed $CWM_{local}$, $CWM_{landscape}$, and $CWM_{global}$ for height, LDMC and SLA.

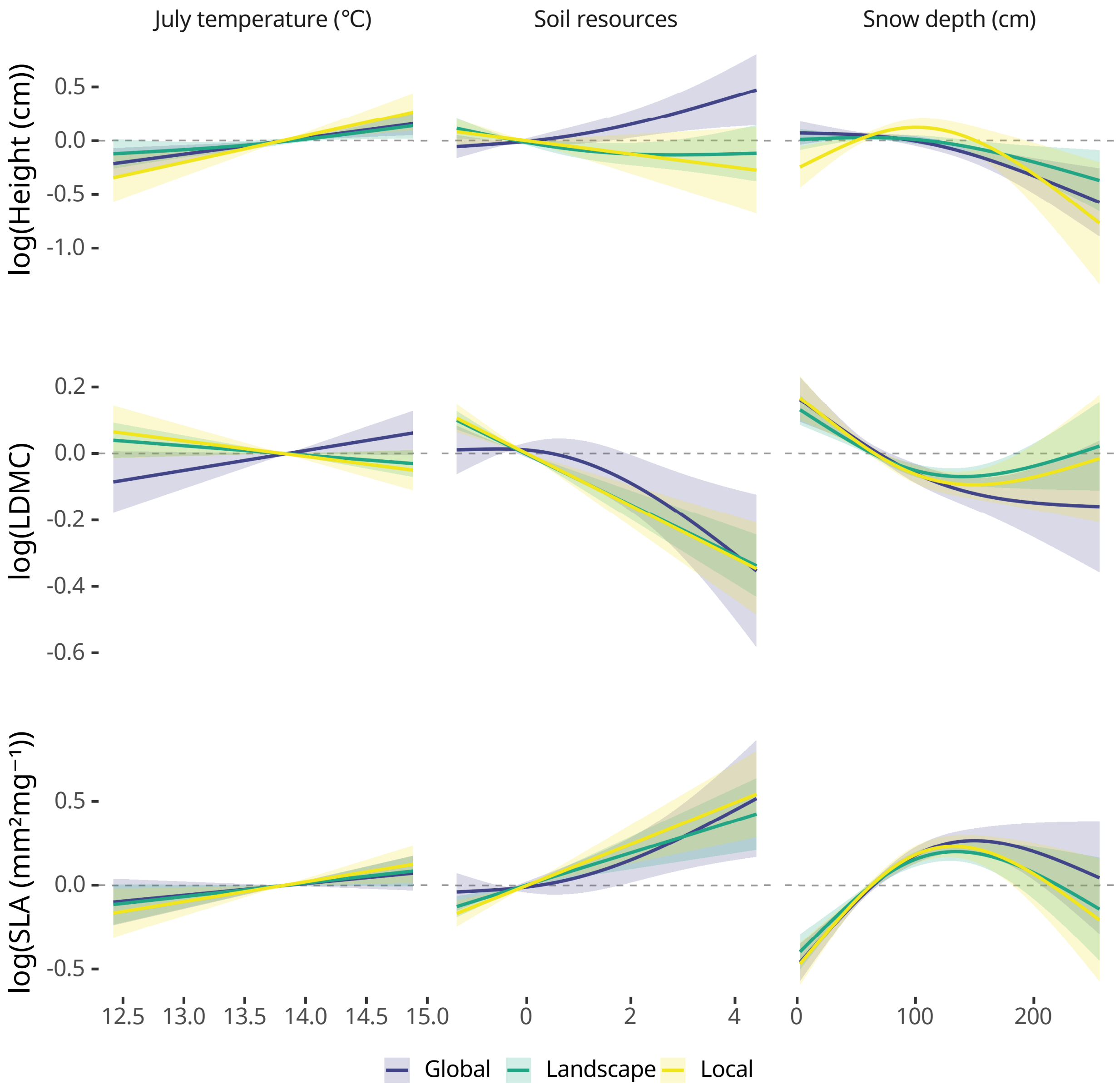

**Figure S6.** Covariation of $\log_e$-transformed $CWM_{local}$, $CWM_{landscape}$, and $CWM_{global}$ between height, LDMC and SLA.

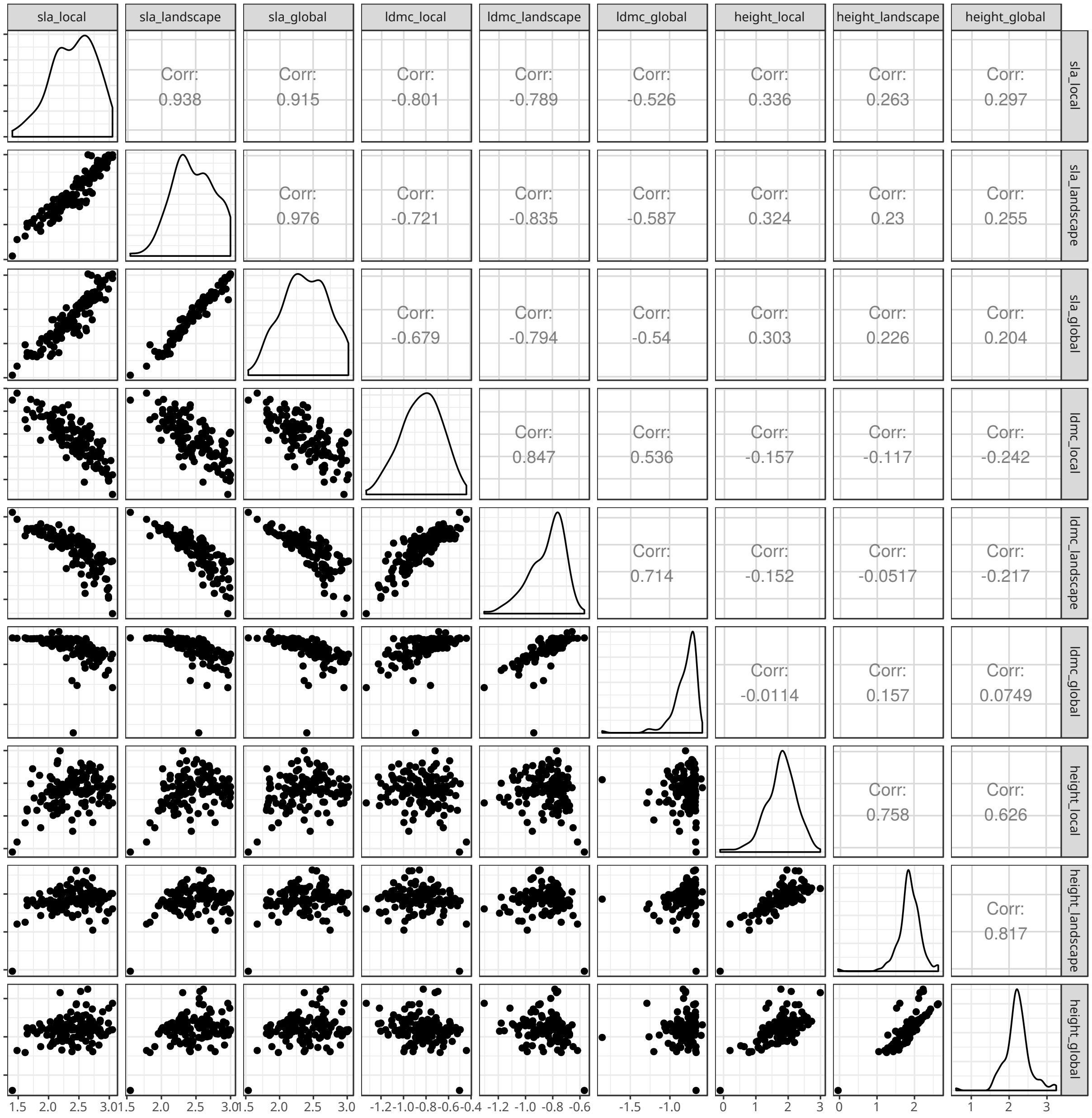

**Figure S7.** Covariation of environmental conditions at the study site.

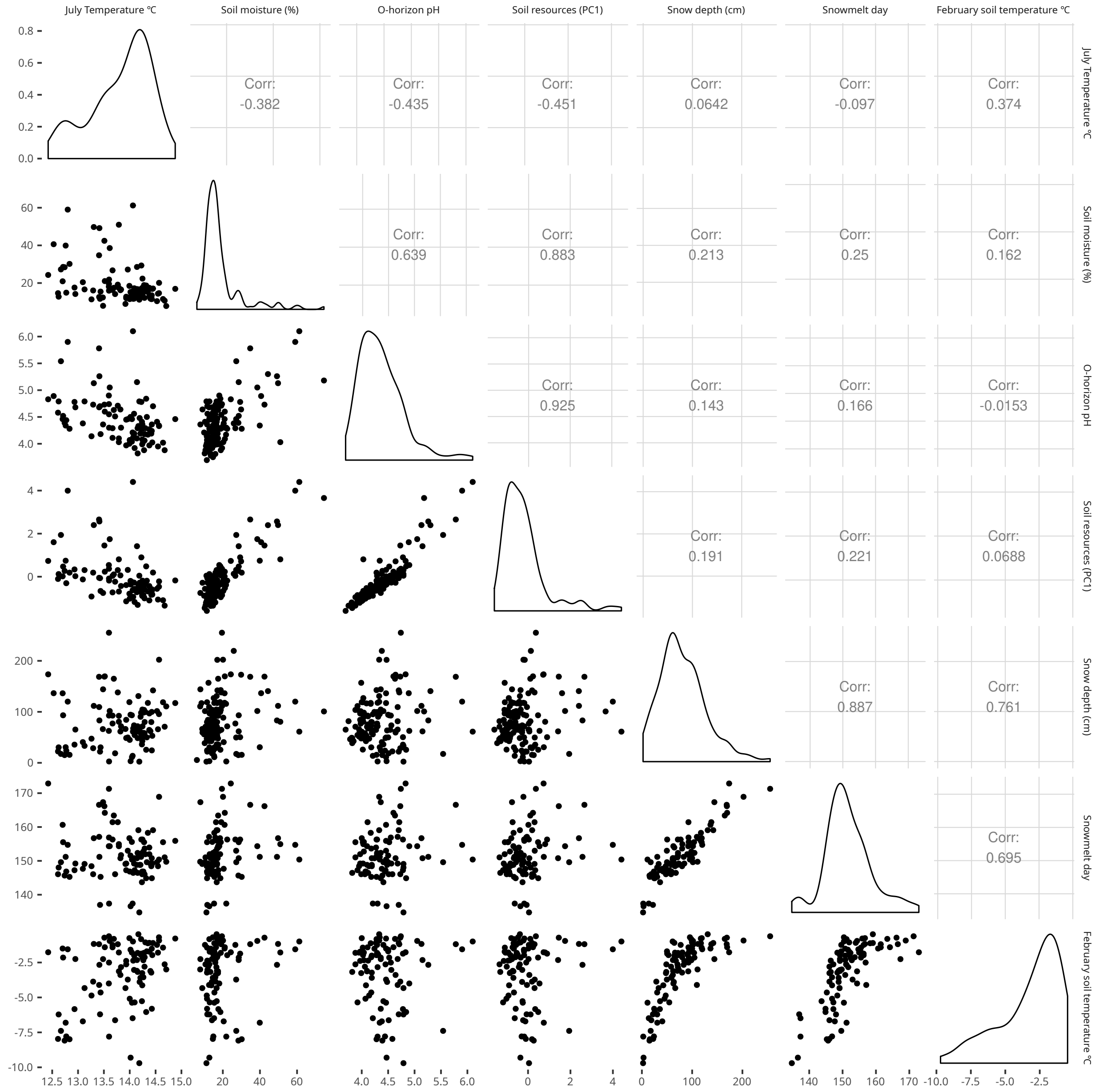
